# The *baseless* mutant links protein phosphatase 2A with basal cell identity in the brown alga *Ectocarpus*

**DOI:** 10.1101/2022.09.10.507423

**Authors:** Olivier Godfroy, Min Zheng, Haiqin Yao, Agnes Henschen, Akira F. Peters, Delphine Scornet, Sebastien Colin, Paolo Ronchi, Katharina Hipp, Chikako Nagasato, Taizo Motomura, J. Mark Cock, Susana M. Coelho

## Abstract

The first mitotic division of the initial cell is a key event in all multicellular organisms and is usually concomitant with the establishment of major developmental axes and cell fates. The brown alga *Ectocarpus* has a haploid-diploid life cycle that involves the development of two multicellular and independent generations, the sporophyte and the gametophyte. Each generation deploys a distinct developmental program autonomously from an initial cell, whose first cell division sets up the future body pattern. Here, we show that mutations in the *BASELESS* (*BAS*) gene result in multiple cellular defects during the first division of the initial cell and subsequently failure to produce basal structures (rhizoids and prostrate filaments) during both generations of the life cycle. Cloning-by-sequencing revealed that *BAS* encodes a type B” regulatory subunit of protein phosphatase 2A, and transcriptomic analysis of early developmental stages uncovered potential effector genes involved in setting up basal cell fate in this organism. The *bas* mutant phenotype is very similar to that observed in the *distag (dis)* mutants, which lack a functional TBCCd1 protein, at both the cellular and morphological levels. The high level of similarity of the *dis* and *bas* mutant phenotypes indicate that TBCCd1 and PP2A are two critical components of the cellular machinery that regulates the division of the initial cell and mediates the establishment of basal cell fate in the developing thallus.

## Introduction

In most animals and plants key events during the first cell division lead to the establishment of one or more major body axes, providing the foundation for the deployment of the subsequent steps of multicellular developmental programs (reviewed in (Radoeva et al., 2019; Rose and Gönczy, 2014). In animals, partitioning defective (PAR) proteins play a key role in establishing the anterior/posterior body axis, whilst a number of pathways are involved in establishing the dorsoventral axis (e.g. Mongera et al., 2019). In the land plant *Arabidopsis*, both the apical/basal axis and apical/basal cell identities are established at the time of the first cell division, and genetic analysis has identified two main pathways involved in this process (reviewed in Bayer et al., 2017; Ueda and Berger, 2019). The first pathway involves SHORT SUSPENSOR (an interleukin-1 receptor-associated kinase/Pelle-like kinase), YODA (a MAP kinase kinase), MPK3 and MPK6 (MAP kinases) and downstream transcription factors, which may include WRKY2 and GROUNDED/RKD4. This pathway may also be influenced maternally through secreted peptide factors such as EMBRYO SURROUNDING FACTOR1 and CLV3/ESR-RELATED8. The second pathway involves auxin and consists of PIN-FORMED 7 (auxin efflux regulator), MONOPTEROS/AUXIN RESPONSE FACTOR 5 (transcription factor) and BODENLOS/INDOLE-3-ACETIC-ACID 12 (auxin response inhibitor). The establishment of zygote polarity is a pre-requisite of the asymmetrical first cell division. This process involves movement of the nucleus and other organelles, enlargement of the vacuole, and reorganization of microtubules (Kimata et al., 2016; Ueda and Berger, 2019).

The mechanisms underlying the onset of early development from an initial cell in multicellular plants and animals are relatively well understood, but research has lagged behind for the third most complex group of multicellular eukaryotes, the brown algae. The brown algae offer an interesting contrast to animals and plants because of their phylogenetic position and the fact that they evolved complex multicellularity independently from those groups. Moreover, in brown algae, the two generations are independent and often very distinct morphologically. The same genome, therefore, regulates the set-up of two independent and distinct developmental programs from two different types of initial cells, opening interesting questions about the molecular control of alternation of generations (Arun et al., 2019; Cock et al., 2014; Coelho et al., 2007; Coelho et al., 2011). Furthermore, gametophytes and sporophytes of brown algae develop from single cells outside the parent organism, indicating that they likely establish polarity in a cell-autonomous manner, without the involvement of factors delivered from the parental tissues, simplifying the study of polarity, axis establishment and pattern formation.

Investigations using the brown alga *Fucus* have shown that asymmetrical first cell division is driven by apical-basal polarity established within the zygotic cell (Brownlee and Bouget, 1998; Goodner and Quatrano, 1993); reviewed in Bogaert et al., 2022). The two daughter cells of this first division divide to produce the apical and basal systems of the alga, the thallus and the rhizoid, respectively (Brownlee and Bouget, 1998). Studies using *Fucus* zygotes have underlined the role of Ca^2+^ asymmetries, mRNA distribution and position-dependent information from the cell wall (involving an unknown diffusible apoplastic factor) in the determination of the fate of the basal and apical systems (Berger et al., 1994; Bogaert et al., 2022; Bouget et al., 1996; Brownlee and Bouget, 1998).

More recently, *Ectocarpus* has emerged as a suitable model to investigate the molecular mechanisms underlying initial cell division and cell fate determination in the brown algae (Bogaert et al., 2022). Advantages of this model include the availability of a range of genetic and genomic tools (Avia et al., 2017; Badis et al., 2021; Bourdareau et al., 2021; Cock et al., 2017; Coelho et al., 2020; Cormier et al., 2017; Umen and Coelho, 2019) but also, importantly, the regularity of the first cell division that characterises the early stages of development of both the gametophyte and sporophyte generations. In this organism, the gametophyte generation exhibits an asymmetrical initial cell division that produces a basal rhizoid cell and an apical cell, the latter further dividing to form the apical system of upright filaments. The upright filaments bear the reproductive structures (plurilocular gametangia, which produce the gametes by mitosis). In contrast, the initial cell of the sporophyte generation undergoes a symmetrical initial cell division to produce two daughter cells with similar fates, both being components of the basal system (Godfroy et al., 2017; Peters et al., 2008)(Figure 1). The apical system of the sporophyte (upright filaments) are produced later, after an extensive system of basal filaments has been established. Reproductive structures (unilocular and plurilocular sporangia) are produced on the upright filaments.

**Figure 1.**
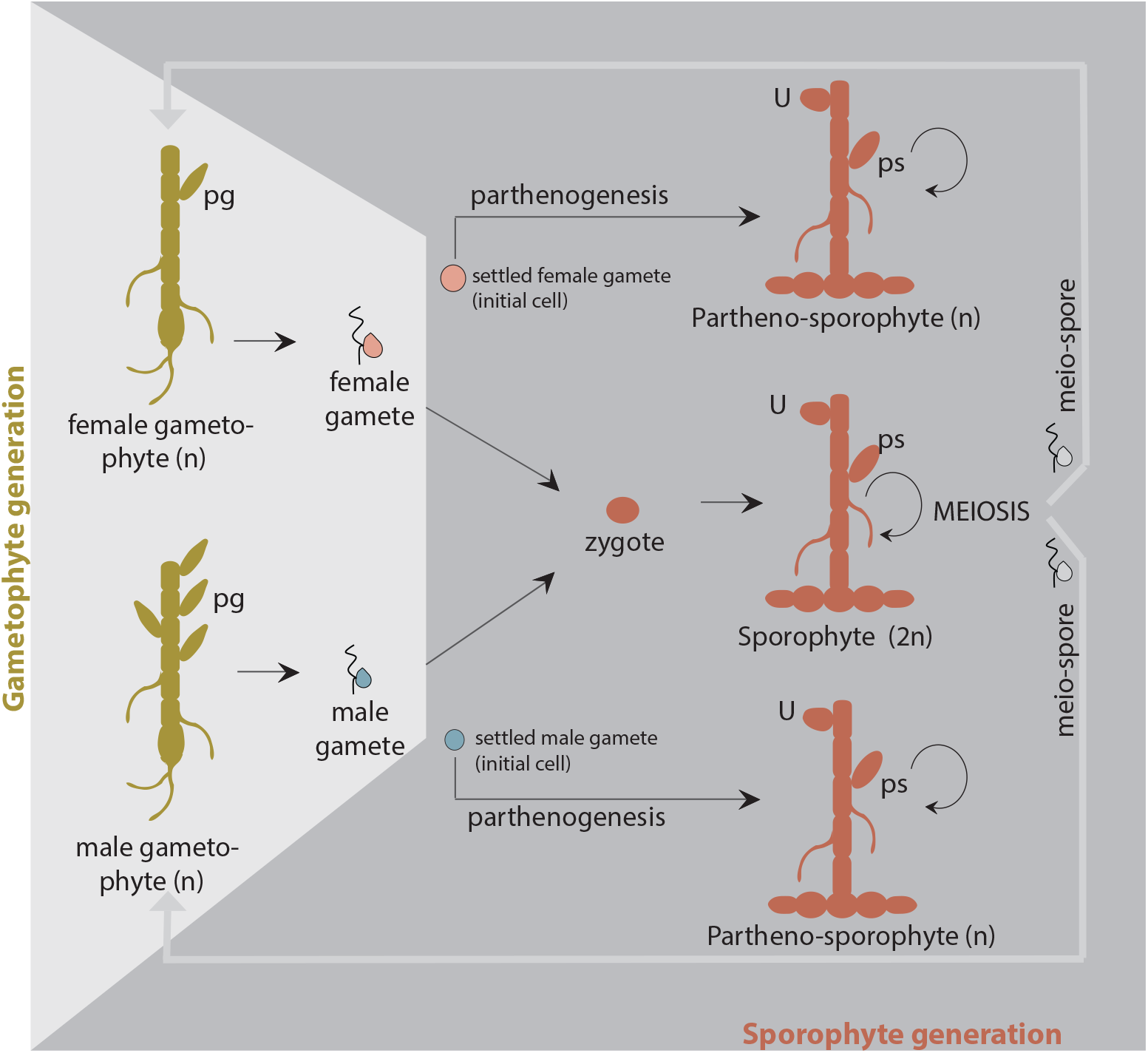
Schematic view of the life cycle of Ectocarpus sp.. Like most brown algae, Ectocarpus has a haploid-diploid life cycle with haploid, genetic sex determination (Coelho and Cock, 2020; Coelho et al., 2020). Male and female gametophytes produce male and female gametes, respectively, which are released into the seawater from plurilocular gametangia (pg). Gamete fusion produces a zygote which will initiate the (diploid) sporophyte generation. However, gametes that do not meet a gamete of the opposite sex settle and lose their flagella, and they may still function as initial cells of the sporophyte generation because they can develop parthenogenically into a (haploid) partheno-sporophyte. Note that there is no visible (morphological) difference between haploid and diploid sporophytes. Meiosis occurs in the sporophyte (in the unilocular sporangia, U), producing several hundred haploid meio-spores. These meio-spores are released into the seawater and develop into male or female gametophytes. Partheno-sporophytes and diploid sporophytes may also be maintained by production of mito-spores in plurilocular sporangia (ps): released mito-spores recycle the partheno-sporophyte generation (circular arrow in the scheme).

Earlier work identified an *Ectocarpus* mutant, *distag (dis)*, with abnormal cellular features during the first cell division, and that was unable to develop basal systems (rhizoids in the gametophyte, prostrate filaments in the sporophyte)(Godfroy et al., 2017). *dis* mutant alleles exhibit a strong phenotype in the initial cell, with disordered microtubule networks, larger cell size, altered Golgi structure and mispositioned nuclei and centrioles (Godfroy et al., 2017). The cell division plane, however, is normal and the cellular defects are only observed during the division of the initial cell, suggesting that *DIS* function may be specific to this cell. *DIS* encodes a Tubulin Binding Cofactor C (TBCC) domain protein of the TBCCd1 class, with conserved roles in animal and plants (André et al., 2013; Feldman and Marshall, 2009).

Here, we report the identification of the *BASELESS* (*BAS*) locus in *Ectocarpus. bas* mutants exhibit phenotypes that closely resemble those of *dis* mutants, including an atypical initial cell division that leads to failure to deploy a basal system in the adult organism, abnormal cellular features such as disorganised microtubule cytoskeleton, loss of bipolar germination and more extensive Golgi apparatus compared with wild-type cells. These phenotypic features are underlie by important modifications in patterns of gene expression during very early stages of development. *BAS* encodes a protein phosphatase 2A regulatory subunit type B” with EF-hand domains. Together, our results are consistent with BAS being involved in a pathway that plays a key role in initial cell division and basal cell fate determination during both the gametophyte and sporophyte generations of the *Ectocarpus* life cycle.

## Results

### *baseless* mutants lack a basal system during both the sporophyte and gametophyte generations

During the *Ectocarpus* gametophyte generation, the two cells derived from the division of the initial cell develop as two germ tubes, and establish a rhizoid (a basal, root-like organ) and an upright filamentous thallus (an apical, shoot-like organ) (Figure 2A, 2B). The first cell division of the sporophyte generation, in contrast, produces two daughter cells with similar morphology and equivalent cell fates (Figure 2C). These two cells then divide to produce a prostrate filament, which branches to establish the basal system and the apical system differentiates later in development, growing up into the medium from the extensive, prostrate basal system. Reproductive structures are produced on the apical system (Figure 2D, 2E).

**Figure 2.**
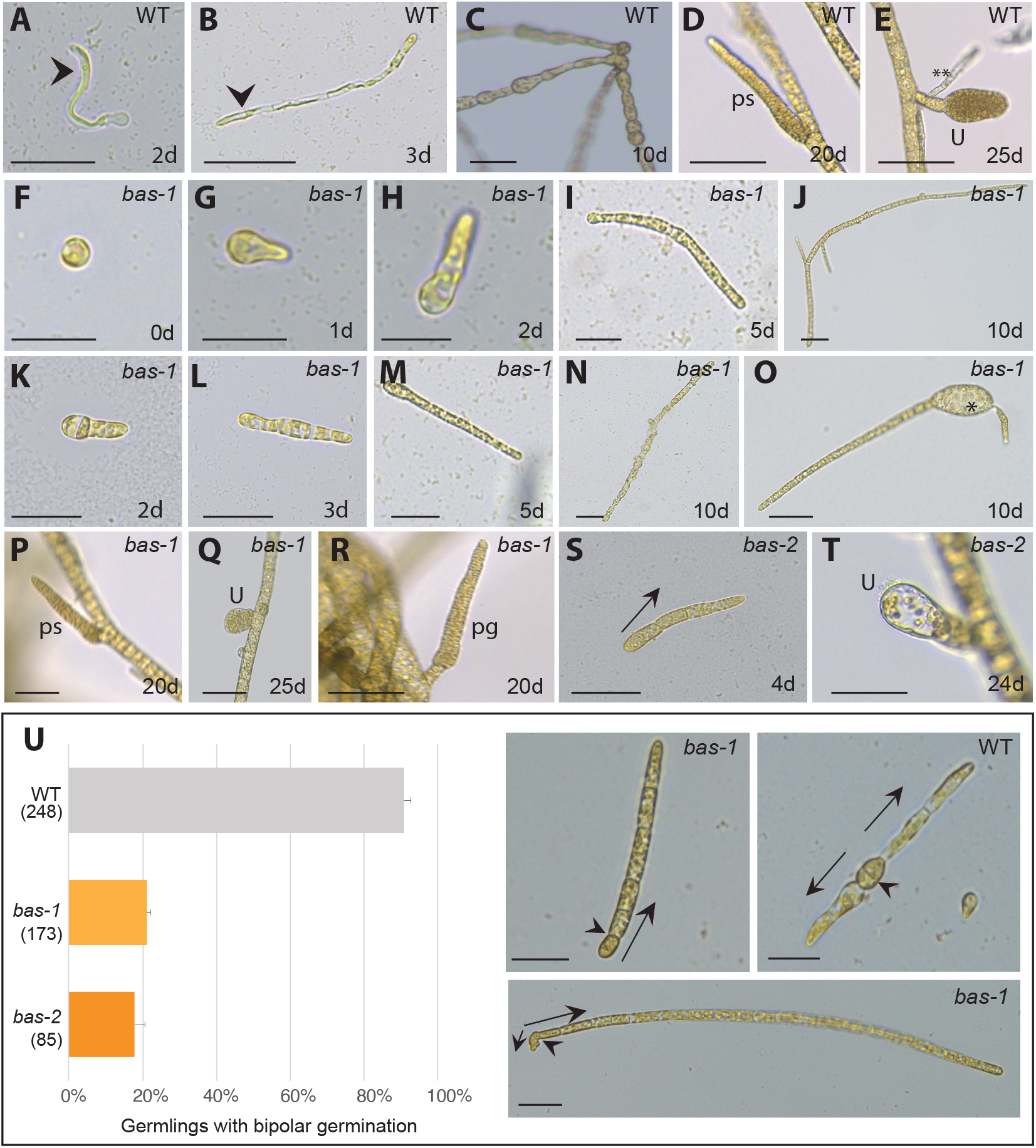
Phenotypes of bas mutants. A) Wild-type gametophyte germling, the arrowhead indicates the rhizoid cell (basal structure). B) Three-day old wild-type gametophyte, the arrowhead indicates the rhizoid. C) Wild-type sporophyte generation composed of round prostrate filaments firmly attached to the substrate. D) Wild-type plurilocular sporangium containing mitotic spores, produced after 20 days in culture. E) Wild-type unilocular sporangium (where meiosis takes place) produced after 25 days in culture. A secondary rhizoid is indicated by asterisks (**). (F-J) Development of the gametophyte generation of the bas-1 mutant. (K-N) Development of the sporophyte generation of bas-1 mutant. O) Occasionally, the mutant strains produced enlarged, abnormal cells (asterisk). P) Plurilocular sporangium on a bas-1 mutant sporophyte. Q) Unilocular sporangium on a bas-1 mutant sporophyte. R) Plurilocular gametangium on a fertile bas-1 gametophyte. S) Initial cell division of a bas-2 gametophyte. T) Aborted unilocular sporangium on a mature bas-2 sporophyte (about 3 weeks after initial cell germination). U) Proportions of 10-day old bas-1 and wild-type germlings that exhibited unipolar germination. Plots represent the mean and SE of five replicate cultures, the total number of germlings scored are indicated in brackets. The photographs are of representative bas-1 and wild-type (WT) germlings, exhibiting uni-(one arrow) or bi-polar (two arrows) germination, respectively. Arrows indicate the direction of germination. Note that, in the bas-1 mutant, following the first cell division, one of the daughter cells continues to divide to produce an upright filament but division of the other daughter cell is arrested. Arrowheads indicate the division plane of the initial cell, which is perpendicular to the growth axis both in the bas mutants and in the wild type. ps, plurilocular sporangium; pg, plurilocular gametangium; u, unilocular sporangium. Scale bars=20 µm.

A screen of a large population of individuals mutagenised by ultraviolet (UV) irradiation identified two mutant strains (Ec800 and Ec801; Table S1) that failed to develop any of the basal structures normally observed during either the gametophyte or the sporophyte generation of wild-type strains (Figure 2). The screen used gametes, which in absence of fertilisation develop into partheno-sporophytes, being thus initial cells of the sporophyte generation (Figure 1) (Coelho et al., 2011; Godfroy et al., 2015; Godfroy et al., 2017; Peters et al., 2008). Initial cells of Ec800 and Ec801 gametophytes immediately developed as apical upright filament cells and no rhizoid cells were produced. Similarly, during the sporophyte generation, neither mutant strain produced the network of prostrate basal filaments typical of the wild-type sporophyte and, instead, the first divisions of the initial cell directly produced an upright filament (Figure 2).

In wild-type *Ectocarpus*, secondary rhizoids, which are analogous to the adventitious roots produced from the stems of some land plants (Atkinson et al., 2014), are produced from apical upright filament cells at a late stage of development (Figure 2E) (Peters et al., 2008). The Ec800 and Ec801 mutants did not produce secondary rhizoids (Figure 2, Figure S1). Hence, production of all basal, attachment structures, both primary and secondary, was abolished in these mutants. Taking into account these phenotypes, the Ec800 and Ec801 mutants were named *baseless-1* (*bas-1*) and *baseless-2* (*bas-2*) respectively.

The establishment of reproductive structures on apical systems in both the gametophyte and sporophyte generations was unaffected in the *bas-1* mutant, which was fully fertile after three weeks in culture (Figure 2F-2S). In the *bas-2* mutant, the formation of the plurilocular sporangia (which contain mito-spores) was unaffected whereas unilocular sporangia (where meiosis takes place) aborted and no functional meio-spores were produced, preventing generation of gametophytes (Figure 2T).

### *bas* mutants exhibit reduced bipolar germination compared with wild-type strains

In wild-type *Ectocarpus*, the majority of the initial cells (91%) exhibited a bipolar pattern of germination, with two germ tubes emerging from opposite poles of the initial cell (Figure 2U; Peters et al., 2008). In contrast, only 21% of the initial cells of *bas-1* partheno-sporophytes exhibited this bipolar pattern of germination, the remaining 79% undergoing unipolar germination (Figure 2U). A proportion of the *bas-1* partheno-sporophytes that exhibited a bipolar germination pattern produced one or more enlarged and abnormally shaped cells at the extremity where the second germ tube would normally emerge, possibly corresponding to an aborted germ tube (Figure 2O). Similar phenotypes were observed for *bas-2* partheno-sporophytes (Table S2).

### Disorganisation of the microtubule cytoskeleton in *bas* mutant initial cells

Mutations at the *DIS* locus strongly affect the organisation of the microtubule cytoskeleton (Godfroy et al., 2017). Because of the similarity between the morphological phenotypes of *bas* and *dis* mutants, we investigated the distribution of the microtubule network during early development of *bas* mutants compared with wild-type germlings (Figure 3A-B). The microtubule cytoskeleton was markedly disorganised in the *bas* mutants, with supernumerary microtubule filaments and a disordered network (Figure 3C). This microtubule phenotype is reminiscent of that of the *dis* mutant (Godfroy et al., 2017). Also, similarly to the *dis* mutant, we did not detect any abnormalities in the positioning of the cell division plane during early development; all *bas* initial cells produced a cell division plane perpendicular to the growing axis (Figure 2U).

**Figure 3.**
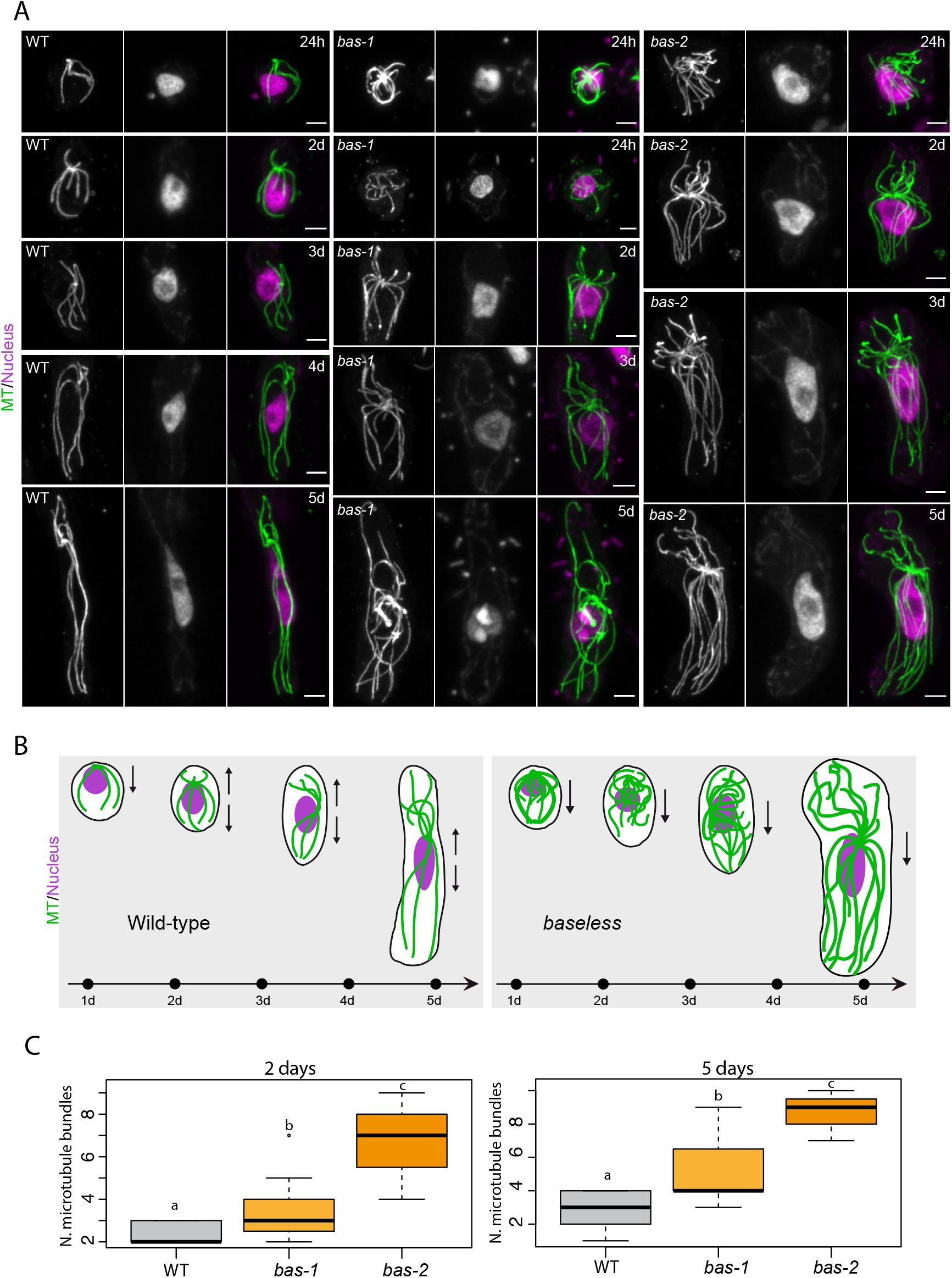
The organization of the microtubule cytoskeleton is affected in bas mutant germinating cells. A) Confocal maximum z-projections showing representative cells of wild type, bas-1, and bas-2 partheno-sporophytes at several stages of early development (24h, 2-5 days). Microtubules (MT) were immune-stained with an anti-tubulin antibody (green). Nuclear DNA was counterstained with DAPI (mauve). Microtubule (MT) bundles were wavy and more abundant in both bas-1 and bas-2 mutant cells compared with the wild-type during the germination of the initial cell. B) Cartoons summarising the stages shown in A) in wild-type and bas mutants. C) Number of microtubule bundles during germination in wild-type (WT), bas-1 and bas-2 mutants at 2 days and 5 days after germination of the initial cell of the sporophyte generation.

### Ultrastructural analysis of *bas* initial cells

Transmission electron microscopy (TEM) and Focused Ion Beam–Scanning Electron Microscopy (FIB-SEM) were used to further characterise the cellular architecture of *bas* mutants. We focused on the early development of the sporophyte generation, i.e., when unfused gametes had started developing into partheno-sporophytes (2-5 cells), which is the stage when conspicuous morphological differences were observed (Figure 2). This is also the stage when *dis* mutants exhibit altered subcellular phenotypes, including significantly more abundant cisternae and more fragmented Golgi compared with wild-type cells (Godfroy et al., 2017).

Morphometric analyses of the subcellular structures in *bas* mutant and wild-type indicated that the Golgi apparatus were about twice as numerous in *bas* as in wild-type at the same developmental stage, but this difference was not significant (Figure 4E; Table S3). We also noticed that the number of cisternae per Golgi tended to be slightly higher in *bas* (Figure 4F, Table S3). Therefore, *bas* exhibited a Golgi defect, but this defect appeared to be less conspicuous than that of *dis* mutants. We did not observe any other organellar defects in *bas* cells, such as abnormal structure or position of the nucleus, centrioles, mitochondria or chloroplasts. Moreover, no visible defect, in particular at the Golgi, could be observed in the gametes prior to their release from the plurilocular gametangia (Figure 4G). This indicates that *bas* cellular defects are detectable only once the initial cell initiates germination.

**Figure 4.**
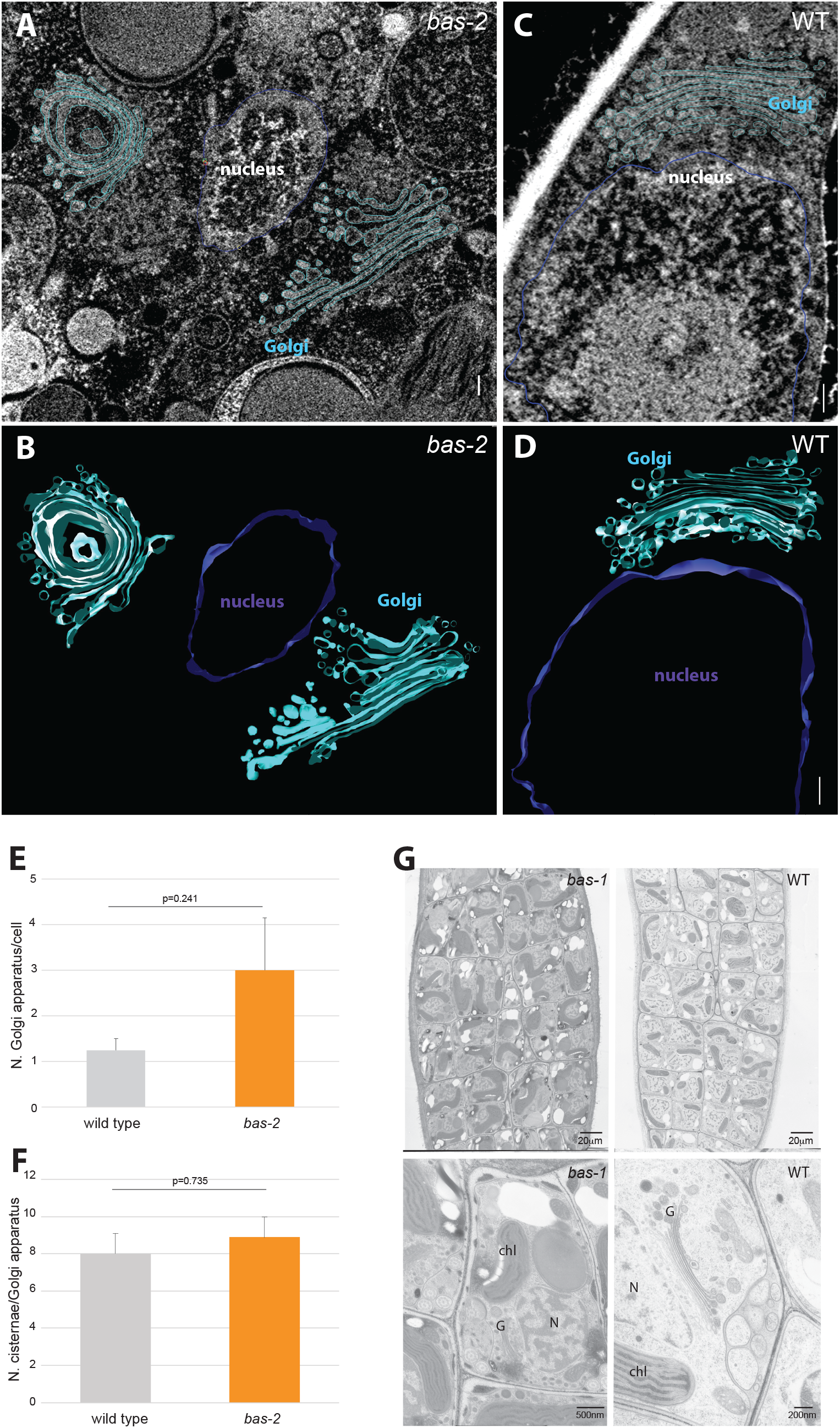
Sub-cellular architecture of wild-type and bas (bas-2) germinating cells. 3D visualization with FIB-SEM (focused ion beam scanning electron microscopy) of representative wild-type (A-B) and bas-2 mutant (C-D) developing sporophytes (2 cell stage), with Golgi (cyan) and nucleus (violet) highlighted. The scale bar represents 200 nm. E) Number of cisternae per Golgi apparatus in wild-type and bas-2 developing sporophytes (2 cell stage). F) Number of Golgi apparatus per cell in wild-type and bas-2 developing sporophytes (2 cell stage). G) Plurilocular gametangia in bas-1 and WT, filled with gametes. Note that no difference in the ultrastructure could be observed in the mutant compared with the wild-type samples prior to the release of the gametes from the plurilocular gametangia. Chl: chloroplast, G: Golgi; N: nucleus.

### Genetic analysis of the BAS gene

A male *bas-1* gametophyte (Ec800) was crossed with a wild-type female gametophyte (strain Ec25; (Table S1 and Figure S2). The resulting sporophyte (Ec805) exhibited a wild-type pattern of development, indicating that the *bas-1* mutation was recessive. A segregating family of 38 partheno-sporophyte individuals derived from this cross consisted of 16 and 22 phenotypically wild-type and mutant individuals, respectively, consistent with a 1:1 segregation ratio and Mendelian inheritance of a single-locus recessive mutation (Chi-square test with Yates correction = 0.4767, p-value = 0.4899; Table S5). The *bas-1* mutation segregated with the phenotype in the 38 individuals used for the phenotype segregation analysis (Table S4).

### *bas-1* and *bas-2* resemble *dis* mutants but are unaffected in the *DIS* gene

The phenotypes of *bas-1* and *bas-2* strongly resembled that of the *dis* mutant (Godfroy et al., 2017). The *dis* mutant also fails to produce any basal structures, during both the sporophyte and gametophyte generations, and lacks secondary rhizoids, again during both generations. Sporophytes resulting from crosses either between the *bas-1* strain Ec800 and strains carrying either the *dis-1* or the *dis-2* allele had wild-type phenotypes (Figure S3 and Table S1), indicating complementation, and therefore that the *DIS* gene was not mutated in the *bas-1* mutants.

### BAS encodes a protein phosphatase 2A type B” regulatory subunit

Genome resequencing identified a candidate locus on chromosome 21 for the location of the *bas-1* and *bas-2* mutations (Figure 5). Whole genome resequencing (WGS) was carried out for the Ec800 (*bas-1*) and Ec801 (*bas-2*) mutant strains and the data compared to the wild-type *Ectocarpus* sp. strain Ec32 reference genome. More than 41,000 putative variants were detected for each strain. Those variants were compared to a list of 567,532 variants called during the analysis of 14 other mutant lines that showed a range of different phenotypes (Table S5) and shared variants were eliminated. This approach allowed the identification of 827 and 769 variants that were unique to the Ec800 and Ec801 mutants, respectively. Quality filtering of those variants (see methods for details) resulted in 118 and 67 putative mutations for the Ec800 and Ec801 strains, respectively, corresponding to one mutation every 1.7 to 3.0 Mb of genome. Of these 185 putative mutations, 26 and 15 were in coding regions (CDS) (22%) in Ec800 and Ec801, respectively. Only one gene (locusID: Ec-21_001770) contained a CDS mutation in both strains. A single nucleotide transition from T to C, at position 2,806,985 was identified in the *bas-1* mutant (strain Ec800) and a G to A transition at position 2,807,321 in the *bas-2* mutant (strain Ec801) (Figure 5). The *bas-1* mutation segregated with the phenotype in the 38 individuals used for the phenotype segregation analysis (Table S6).

**Figure 5.**
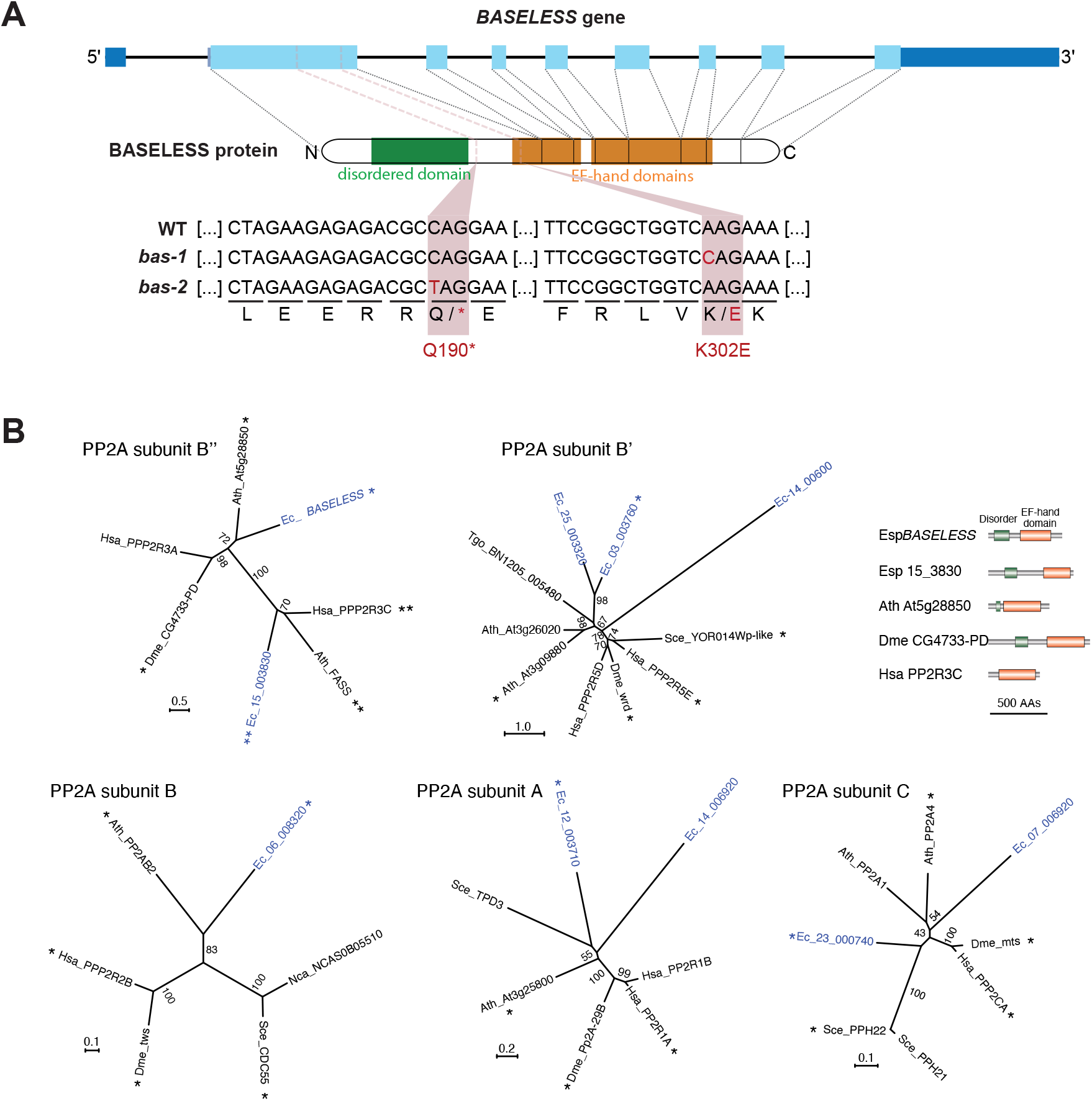
Identification of mutations in the BAS gene and identification of protein phosphate 2A subunits in Ectocarpus. A) Diagram showing the domain structure of the BAS gene, indicating the positions of the bas-1 and bas-2 mutations. The point mutation in exon 2 in bas-1 results a lysine (K) being replaced with a glutamate (E) residue, whereas the point mutation in bas-2 results in the introduction of a stop codon into the coding region of the gene (represented by an asterisk). Blue boxes indicate exons, with dark-blue representing untranslated regions and light-blue the coding region. B) Ectocarpus protein phosphatase 2A subunits. Unrooted maximum likelihood trees of PPA2A subunits (LG+G model). Only bootstrap (1000 repetitions) values of >50 are shown. Ectocarpus proteins are shown in blue. Asterisks and double asterisks indicate best species-to-species reciprocal Blastp matches with the corresponding Ectocarpus protein. The Ectocarpus genome does not encode an orthologue of PP2A subunit B”‘/striatin. The domain structures of five PP2A subunit B” proteins are shown with the EF-hand domains in brown and disordered domains in green. AA, amino acid; Esp, Ectocarpus sp.; Tgo, Toxoplasma gondii; Ath, Arabidopsis thaliana; Hsa, Homo sapiens; Dme, Drosophila melanogaster; Sce, Sccharomyces cerevisiae; Nca, Naumovozyma castelli. Ectocarpus locusIDs are abbreviated as in the following example: Esp_14_3830, Ectocarpus sp. Ec-15_003830.

The Ec-21_001770 gene encodes a protein of 646 amino-acids similar to protein phosphatase 2A regulatory subunit type B” proteins. This polypeptide contains three predicted functional domains: a disordered region between positions 50 to 185 and two EF-hand domains at positions 280 to 370 and 380 to 550. The *bas-1* mutation affects the first EF-hand, replacing a positively charged lysine residue with a negatively charged glutamic acid (K302E). This modification of electric charge may disrupt domain folding and/or function at least for the first EF-hand. It is possible that the *bas-1* mutation leads to the production of a protein that is partially active. The *bas-2* mutant carries a non-sense mutation that creates a premature stop codon at position 190 of the protein. This mutation is predicted to result in the production of a truncated protein that lacks both EF-hand domains (Figure 5A).

*BAS* is predicted to encode a protein phosphatase 2A regulatory B” subunit. PP2A phosphatases are protein complexes usually composed of three subunits, a catalytic C subunit, a scaffolding A subunit and a regulatory B subunit (Wlodarchak and Xing, 2016). Most species have multiple forms of each subunit and there are four distinct classes of the B subunit (B/B55/PR55, B*’*/B56/PR61, B*’’*/PR72/PR130 and B*’’’*/Striatin), which are unrelated at the sequence level. An analysis of the *Ectocarpus* sp. genome revealed that it encodes B, B’ and B” subunits, but not B’”/Striatin (Figure 5B). The BAS protein is predicted to belong to the B” class, homologous to the PR130/PR72 human protein (Figure 5B).

### *BAS* expression during *Ectocarpus* life cycle and in other developmental mutantsz

RNA-seq data (Bourdareau, 2018; Coelho et al., 2011; Godfroy et al., 2017; Macaisne et al., 2017) were analysed to investigate *BAS* gene expression during the *Ectocarpus* life cycle. *BAS* transcripts were detected throughout development, during both the gametophyte and sporophyte generations (Figure 6A). The *BAS* transcript was most abundant at the gamete stage (after the release from plurilocular gametangia) and had decreased in abundance about one week after germination, at the 2-5 cell stage (Figure 6A; Tables S7). This pattern of expression is consistent with a role of BAS in the early divisions of the initial cells of the partheno-sporophyte generation provided that the transcript and/or protein persists in the initial cell during the first cell division. Interestingly, during the life cycle, the abundance of the *BAS* transcript was inversely related to the abundance of the *DIS* transcript, which was at a very low level in gametes but increased in abundance at later stages of development (Figure 6, Table S7, S8).

**Figure 6.**
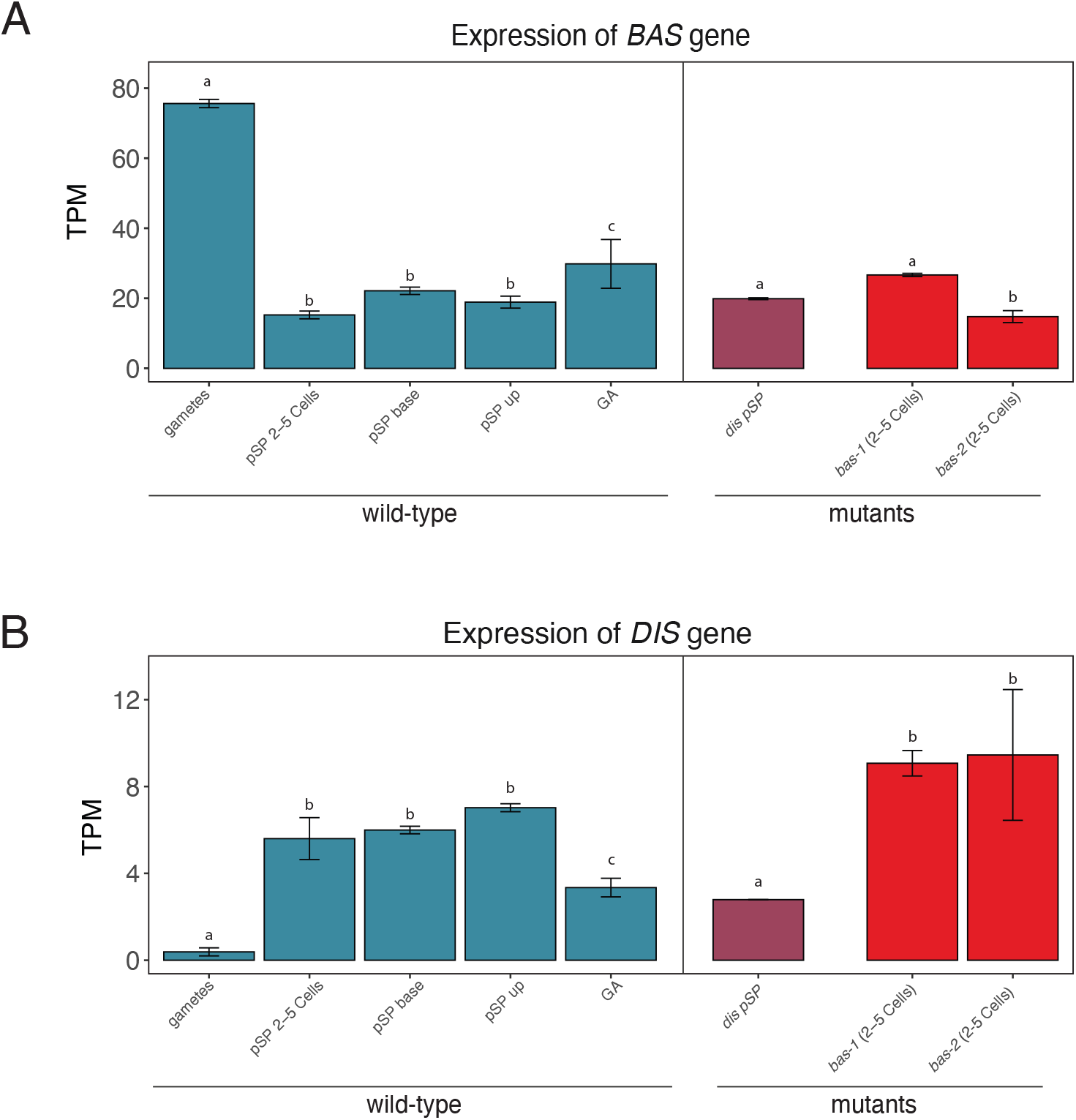
Abundance of the BAS and DIS transcripts (measured as transcripts per million, TPM) during the life cycle of Ectocarpus and in developmental mutants. A) BAS transcript abundance during several developmental stages of wild-type Ectocarpus and in dis and bas mutant strains. Note the high abundance of the BAS transcript in gametes. GA: gametophyte, pSP: partheno-sporophyte. Significant differences in expression (Tukey test) are indicated as different letters above the plots, and detailed statistics are presented in Table S14. B) Abundance of the DIS transcript in wild-type samples compared with developmental mutants. pSP: partheno-sporophyte; GA: gametophyte; up, upright filaments of the pSP; base, basal system of the pSP; (2-5 Cells), early development (2-5 cell stage) of the partheno-sporophyte.

The similar phenotypes of *dis* and *bas* mutants suggest that the products of the two genes may play roles in common cellular processes. We investigated the expression of *DIS* in a *bas* background, and, conversely, the expression of *BAS* in a *dis* background. We noticed that in a *bas* background, *DIS* expression in early stage (2-5 cell) was significantly higher (p-value=4.13E-02 and p-value=7.66E-02 respectively for *bas-1* and *bas-2* comparison with wild-type at the 2-5 cell stage, Figure 6B; Table S7), whereas no difference was observed in the levels of expression of BAS in absence of *dis* gene product. Taken together, these analyses suggest that *BAS* expression levels are not affected by *DIS*, but, conversely, *DIS* gene expression is disturbed by mutations at the *BAS* locus. This observation, however, is unlikely to explain the phenotypic similarity between *bas* and *dis* mutants.

### Analysis of the *bas* transcriptome

To further characterize the *bas* phenotype, an RNA-seq approach was employed to study gene expression in the *bas-1* and *bas-2* mutants compared with wild-type. We focused on the 2-5 cell stage during gamete germination because the *bas* phenotype was prominent during the early stages of development. Given the subtle differences in phenotype between *bas-1* and *bas-2* (in particular the meiotic defect in *bas-2*) we also focused on comparing *bas-1* and *bas-2* transcriptional patterns.

Overall, gene expression patterns in *bas-1* and *bas-2* were comparable, and both were different from the wild-type samples (Figure S4-S5; Tables S7-S9). However, *bas-2* exhibited more differences compared with wild-type samples (Figure S4-S5): 40% of the transcriptome was differentially regulated in the comparison *bas-2* versus wild-type, whereas 20% of the genes was differentially expressed (DE) between *bas-1* and wild-type (Figure S6, Table S8, see Material and Methods for thresholds). Most genes were expressed under all conditions, only about 4% of the genes were not expressed in any of the samples.

Differentially expressed (DE) genes exhibited very similar patterns of up and down regulation in the two *bas* mutants compared to wild-type. Only 11 genes exhibited divergent expression patterns, i.e., up-regulated in *bas*-1 and down-regulated in *bas*-2, and six genes were up-regulated in *bas*-2 and down-regulated in *bas*-1. The vast majority of the DE genes in *bas*-1 vs WT were also differentially expressed in *bas*-2 vs wild-type comparison (76.69% of the down-regulated genes and 76.73% of the up-regulated genes) (Figure 7A, Table S9).

**Figure 7.**
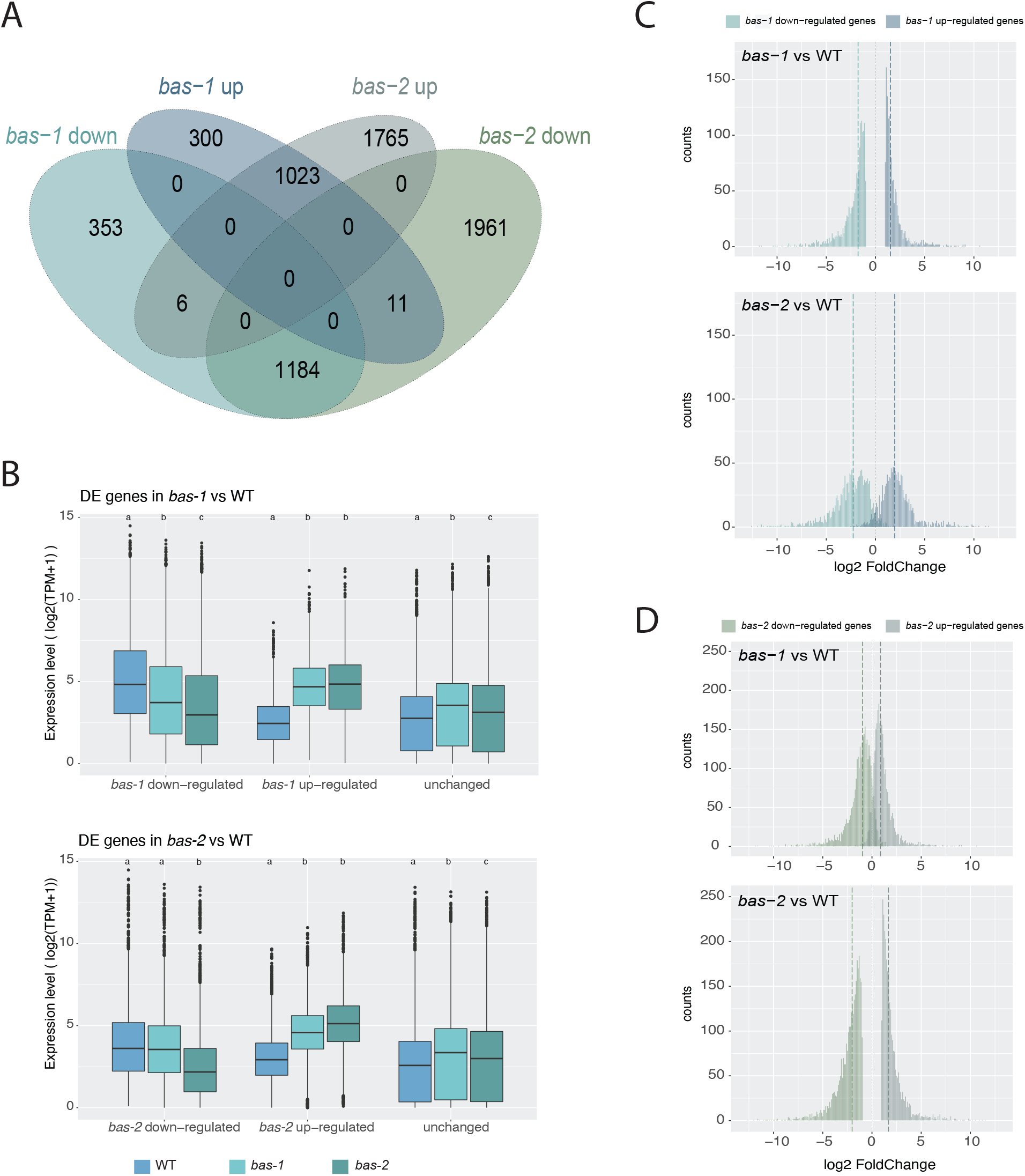
Differential gene expression analysis. A) Venn diagram of intersects of DE gene sets in bas mutants compare to wild-type. B) Boxplot representation of the distribution of gene expression levels (in log2 of TPM values +1) of the DE gene sets from comparisons of either bas-1 or bas-2 with wild-type. For each gene category, significant differences in expression level across strains, according to the Wilcoxon test, are indicated as different letters above the plots (details in Table S15). C-D) Histogram representation of the distribution of fold changes of the DE gene sets from comparisons of bas-1 or bas-2 with WT; vertical dashed bars indicate the medians of the distributions. DE genes in bas-1 compared to wild-type (C); DE genes in bas-2 compared to wild-type (D).

Down-regulated genes in *bas-1* compared to wild-type had, globally, lower expression levels in *bas-2*; and up-regulated genes in *bas-1* were slightly more expressed in *bas-2* (Figure 7B). Conversely, down-regulated genes in *bas-2* compared to WT had expression levels in *bas-1* similar to wild-type samples; while up-regulated genes in *bas-2* had intermediate expression level compare to wild-type and *bas-2* (Figure 7B, Table S9). Comparison of the fold-change distribution of the DE genes from *bas-1* vs wild-type with those of the DE genes from *bas-2* vs wild-type showed that DE genes exhibited greater fold changes in *bas*-*2* than in *bas-1* (Figure 7C, D).

Overall, these observations indicate similar changes in transcriptome of both *bas* mutants, with a stronger effect in *bas-2* than in *bas-1* compared to wild-type. Those results are coherent with the predicted effects of the two mutations on the *BAS* protein. The *bas-1* mutation may result in the production of a protein partially active whereas the *bas-2* mutation leads to a truncated protein with probably no activity.

GO term enrichment analyses of DE genes revealed that functions related to photosynthesis and metabolism were down regulated in both *bas* mutants whereas upregulated genes in mutants were associated with intracellular protein transport transcription and protein synthesis (Figure S7, Table S10). It is interesting to note that some of those functions can be linked with the Golgi apparatus, which appears to be affected by the *bas* mutation (see above). Using the HECTAR predictor (Gschloessl et al., 2008), we examined the enrichment of DE genes in particular sub-cellular localizations. In coherence with GO term enrichment analyses, we observed an enrichment in “chloroplast” localization of the down-regulated genes in both *bas* mutants. We also observed a slight but strongly significant enrichment in proteins with a “signal peptide” suggesting an impact of the *bas* mutation on the production of secreted proteins (Table S11), which, again, may be consistent with defects observed in *bas* mutants at the level of the Golgi. Finally, we noted that 13 out of the 26 predicted BASIC LEUCINE-ZIPPER (bZIP) transcription factors in *Ectocarpus* are differentially expressed in *bas-2* mutant compared to wild-type (Table S9).

We identified 26 and 41 genes that were exclusively expressed in the *bas-1* and *bas-2* respectively compared with the wild-type (i.e., genes that had TMP=0 in wild-type samples). Seventeen genes were silenced in the *bas-1* and 29 in *bas-2* compared with wild-type (Table S12 and S13, see Material and Methods for thresholds). Most of those genes have unknown functions and, due to the low numbers of genes, no significant functional enrichment could be established. However, it is worth noting several transcription factors, which may be potential effector genes: such as Ec-14_003940 and Ec-16_000350, which are silenced in *bas-1* and/or *bas-2* and Ec-00_004890, which is activated in both mutants compared with the wild-type; and numerous genes associated with “Cellular regulation and signalling” or “Membrane function and transport”. Looking at the putative localisation of those proteins, we observed that about 30% of the genes specifically silenced in *bas-1* and/or *bas-2* have “signal peptide” or “signal anchor” prediction, suggested a membrane or cell-wall localization, whereas only 16% of the total predicted proteins in the genome of *Ectocarpus* present this signature (Table S12). This two-fold enrichment is even higher than the “signal peptide” enrichment found in down-regulated genes (Table S11).

## Discussion

### The *BAS* gene is involved in apical-basal axis formation in *Ectocarpus*

The two *Ectocarpus* mutant alleles identified in this study, *bas-1* and *bas-2*, lack basal structures during both the gametophyte and the sporophyte generations of the life cycle. Analysis of the initial cells of the *bas* mutant showed that the morphological phenotype was associated with several cellular anomalies during germination and the first cell division, including disorganisation of the microtubule network, an increase in the number of microtubule bundles and Golgi apparatus and unipolar, rather than bi-polar, germination patterns. These observations highlight a key role for the BAS protein during the development of *Ectocarpus*. In particular, the BAS protein appears to operate during key cell divisions during development: the (first) initial cell division and, at later stages of development, during meiosis. However, no cellular defect was observed prior to initial cell divisions, suggesting that BAS operates only after release of the initial cells from the reproductive structures. The meiotic defect observed in the *bas-2* mutant suggests that the BAS protein may have also a role during meiotic cell division. The absence of this meiotic defect in the *bas-1* may be explained by the production of a protein with sufficient activity in *bas-1* to ensure its role during meiosis but not during the first cell division. Higher penetrance of the *bas-2* mutation is also indicated by the greater proportion of differentially expressed genes in *bas-2* compared to *bas-1*. Indeed, about 40% of the genome was significantly mis-regulated in *bas-2* mutant during early stages of development, demonstrating that mutations at the *BAS* locus can lead to broad, large-scale modifications to the transcriptome. It is interesting to note that half of the bZIP transcription factors of *Ectocarpus* are differentially expressed in *bas-2* compared to wild-type. This observation suggests that BAS may part of a pathway involving additional regulatory proteins that drives the establishment of apical-basal axis during the early development of *Ectocarpus*.

### *BAS* Encodes a PP2A protein with roles in cellular organization and development in animals and plants

*Ectocarpus BAS* is predicted to encode a PP2A phosphatase regulatory B” subunit. In animals, PP2A phosphatases are involved in diverse cellular processes and constitute a major component of cellular serine/threonine phosphatase activity, dephosphorylating several hundred cellular substrates (Reynhout and Janssens, 2019; Wlodarchak and Xing, 2016). PP2A has been implicated in the reorganization of several cellular structures, playing key roles in nuclear envelope breakdown during mitosis, and chromosome segregation via effects on assembly of the mitotic spindle and attachment of cytoplasmic microtubules to kinetochores. It is also involved in rearrangement of endoplasmic reticulum and the Golgi apparatus (reviewed in (Wlodarchak and Xing, 2016; Wurzenberger and Gerlich, 2011). In particular, the PP2A B’’ subunit PR130 associates with CG-NAP that localizes to centrosomes and the Golgi apparatus (Takahashi et al., 1999), and restacking of newly formed Golgi cisternae requires dephosphorylation of Golgi stacking proteins by PP2A (Tang et al., 2008). More broadly, the animal PP2A B’’ subunit is critical for cell-cell communication, cell adhesion, migration, proliferation and differentiation during animal development (Creyghton et al., 2005; Zwaenepoel et al., 2010). Altogether, these observations link animal PP2A to roles in cell division, subcellular features and developmental pattern formation. In the multicellular brown alga *Ectocarpus*, our results are consistent with a similar role for PP2A in subcellular organisation, cell division and cell fate determination. Our results therefore provide an example of protein functional conservation across eukaryotes that evolved multicellularity independently, suggesting that the role of PP2A in multicellular development has likely been preserved across very divergent lineages.

Consistent with a conserved role for PP2A in development across multicellular eukaryotes, the *Arabidopsis* B’’-δ/ε subunit of PP2A (At5g28900/At5g28850) interacts with BASIC LEUCINE-ZIPPER (bZIP) transcription factors, and is implicated in leaf and root development as well as mechanical stress response (Tsugama et al., 2019; Van Leene et al., 2016). The PP2A regulatory B” subunit FASS/TON2, is essential for the reorganisation of cortical microtubular arrays into a dense preprophase band preceding cell division. FASS-containing PP2A complexes are targeted to microtubules through an association with TONNEAU1 (TON1) and TON1-recruiting motif protein (TRM) (Spinner et al., 2010). PP2A interacts with and dephosphorylates KATANIN, a evolutionarily conserved microtubule-severing enzyme, to promote the formation of circumferential cortical microtubule arrays in *Arabidopsis* (Ren et al., 2022). The *Ectocarpus* BAS protein is related to the *Arabidopsis* FASS/TON2 protein but is orthologous to the AtB”-δ/ε protein mentioned above (At5g28900/At5g28850). The microtubule phenotype we describe here, where PP2A B’’ disruption in brown algae is associated with microtubule disorganisation, may further underline a conserved role for PP2A across plants and brown algae.

Golgi vesicle transport has been shown to play an important role in the establishment of the *Fucus* zygote polar axis prior to the first cell division (reviewed in Bogaert et al., 2022; Shaw and Quatrano, 1996)). It has been proposed that the selective targeting of Golgi vesicles to the plasma membrane locally modifies the cell wall, participating in the establishment of the asymmetry of the cell wall required for rhizoid differentiation (Bogaert et al., 2013; Bogaert et al., 2022; Goodner and Quatrano, 1993). In this model, the cell wall plays a key role in the establishment of the initial cell asymmetry, which is interesting with regard to the dramatic changes in expression of numerous genes with putative “Membrane and transport” functions in *bas* mutants.

In animals, the B’’ class of PP2A B subunit is thought to be involved in Ca^2+^ signalling through the presence of multiple EF-hand domains (Xu et al., 2010). Note that two EF-hand domains are predicted in the BAS protein, and these EF-hands are absent or non-functional in *bas* mutants, suggesting that Ca^2+^ binding may be disturbed. In the brown alga *Fucus*, Ca^2+^ gradients have been shown to have a crucial function in zygote polarization and the establishment of the apical/basal axis during first cell division (reviewed in (Bogaert et al., 2022; Brownlee and Bouget, 1998). In particular, cytosolic free Ca^2+^ accumulates on the side of the growing (basal) rhizoid, directly linking intracellular Ca^2+^signaling and acquisition of basal cell identity. In this context, we can speculate that the BAS protein may be a potential sensor of a pre-germination Ca^2+^ signal.

### BAS and DIS may act in concert to mediate cell fate determination during the first cell division

Similar morphological and cellular phenotypes were observed for *bas* and *dis* mutant strains suggesting that *BAS* and *DIS* may be involved in similar cellular processes. However, some differences between the mutants are also apparent, such as the less marked Golgi fragmentation and no effect on nuclear positioning in early stages of development in *bas*, and also a meiotic defect was only observed in *bas-2*.

*DIS* is predicted to encode a TBCCd1 protein. This protein shares similarity with TBCC, which is a component of the complex (TBCA to TBCE) that mediates dimerization of α and β tubulin subunits to form microtubules (Nithianantham et al., 2015; Tian et al., 1996). However, TBCCd1 lacks a conserved arginine residue that is essential for TBCC activity and is unable to complement TBCC in yeast indicating that the two proteins may have different biochemical functions (Goncalves et al., 2010). TBCCd1 has been localized to both the centrosome and the Golgi in humans, *Chlamydomonas*, and trypanosomes and there is evidence that TBCCd1 plays important roles in positioning organelles within the cells of these diverse organisms (André et al., 2013; Feldman and Marshall, 2009; Goncalves et al., 2010). However, the molecular mechanisms underlying these cellular phenotypes are unclear and they may not involve direct effects on microtubule assembly (Goncalves et al., 2010).

Interestingly, analysis of human protein phosphatase interactions revealed that TBCCd1 is a partner of PP2A regulatory subunit B’’ (Huttlin et al., 2015; Huttlin et al., 2017; Yadav et al., 2017). If this interaction also occurs in brown algae, it would provide a mechanism whereby DIS and BAS could act within the same pathway involved in cell fate determination, with BAS regulating the DIS protein. Further analysis of the biochemical functions of BAS and DIS will be necessary to test this hypothesis.

To summarise, both TBCCd1 and PP2A have been linked to cytoskeleton and Golgi function and both proteins have been shown to play important roles in the regulation of cellular architecture in diverse eukaryotic systems. These observations are consistent with the pleiotropic cellular phenotypes of both the *dis* and *bas* mutants. We suggest that the observed morphological and cell fate (loss of basal cells) phenotypes of the *bas* and *dis* mutants are a consequence of cellular defects during the first cell division, perhaps through disruption of the distribution of hypothetical cell-fate-determining factors during this critical step of development (see model proposed by (Godfroy et al., 2017). Combining information about the *Ectocarpus* DIS and BAS proteins with observations of Ca^2+^ waves in the *Fucus* embryo, we can speculate that BAS may be involved in sensing an intracellular Ca^2+^ signal which participates to the distribution of a cell-fate determining factor through reorganization of cytoskeleton and Golgi function involving DIS.

## Methods

### UV Mutagenesis and isolation of mutant strains

Strain cultivation, genetic crosses, raising of sporophytes from zygotes, and isolation of meiotic families were performed as described previously (Coelho et al., 2012a; Coelho et al., 2012b; Coelho et al., 2020; Godfroy et al., 2017). *Ectocarpus* sp. (species 7, Montecinos et al., 2017) gametes are able to develop parthenogenically to produce haploid partheno-sporophytes, which are identical morphologically to the sporophytes that develop from diploid zygotes (Bogaert et al., 2022; Bothwell et al., 2010; Peters et al., 2008). This phenomenon was exploited to screen directly, in a haploid population, for mutants affected in early sporophyte development. UV mutagenesis of gametes was performed as described previously (Coelho et al., 2011; Godfroy et al., 2015; Godfroy et al., 2017) and mutant partheno-sporophytes lacking basal structures were identified by visual screening under a light microscope. Table S1 describes the strains used in this study.

### Genetic analysis of *bas* mutants

Genetic crosses were performed as in (Coelho et al., 2012a). The *bas-1* mutant (Ec800) was crossed with the outcrossing line Ec568 to generate a segregating population of 38 individuals. Each of the 38 individuals was derived from a different unilocular sporangium (each unilocular sporangium contains 50–100 meio-spores, derived from a single meiosis followed by at least five mitotic divisions). The meio-spores germinated to produce gametophytes, which were isolated and allowed to produce gametes which germinated parthenogenically. The resulting partheno-sporophytes were then observed under a light microscope to determine whether they exhibited the *bas* phenotype. The presence of the *bas-1* mutation was determined by Sanger sequencing of PCR products (Forward: TGACGAATGATGCTAAACTGGA, Reverse: GACAACGGAGCAGACGAAC) for each of the 38 individuals. The *bas-2* mutant (Ec801) was not usable for crosses because it did not form functional unilocular sporangia, which are required for gametophyte production.

### Identification of candidate mutations

Genomic DNA from Ec800 and Ec801 strains was sequenced on an Illumina HiSeq4000 platform (1/12^th^ lane; 2×150nt paired-end; 8.50 and 7.95 Gbp of data, respectively; Fasteris, Switzerland). After quality cleaning using Trimmomatic (Bolger et al., 2014), the reads were mapped onto the *Ectocarpus* sp. reference genome (Cormier et al., 2017) using Bowtie2 (Langmead and Salzberg, 2012). Coverage depth and breadth were, respectively, 34x and 96.83% for Ec800 and 32x and 96.81% for Ec801. Variants were called and counted using bcftools mpileup (http://samtools.github.io/). These variants were compared with a list of variants identified in genome sequence data for 14 other *Ectocarpus* mutant lines in order to remove false positive mutations due, for example, to errors in the reference genome sequence.

Variants unique to strains Ec800 and Ec801 were quality filtered based on coverage depth (±50% of the genome mean), mapping quality (>20), variant quality (>50), variant frequency (>0.9) and variant support in both sequencing directions. A custom python script allowed the identification of variants in coding regions (CDS) and the effect of each CDS mutation on the predicted protein was accessed manually (Table S5). A scheme of the approach is depicted in Figure S7.

### Immunostaining

*Ectocarpus* samples were processed as described (Coelho et al., 2012c) using a protocol adapted from (Bisgrove and Kropf, 1998). Briefly, *Ectocarpus* cells were settled on cover slips and at appropriate times after settlement were rapidly frozen in liquid nitrogen and fixed in 2.5% glutaraldehyde and 3.2% paraformaldehyde for 1 h, then washed in PBS and treated with 5% triton overnight. Samples were then rinsed in PBS and 100 mM NaBH^4^ was added for 4 h. Cell walls were degraded with cellulase (1% w/v) and hemicellulase (4% w/v) for 1 h, and the preparation was then rinsed with PBS and blocked overnight in 2.5% non-fat dry milk in PBS. Samples were treated with an anti-tubulin antibody (1/200th, DM1A; Sigma-Aldrich) at 20°C overnight and then treated with the secondary antibody (AlexaFluor 488-conjugated goat anti-mouse IgG; Sigma-Aldrich; 1:1000 in PBS) at 20°C overnight. The preparation was rinsed with PBS and blocked overnight in 2.5% non-fat dry milk in PBS and then treated with an anti-centrin antibody (1/1000th anticentrin 1 ab11257; Abcam) at 20°C overnight, followed by the secondary antibody (1/1000th AlexaFluor 555-conjugated goat anti-rabbit IgG; Sigma-Aldrich) for 8 h. Samples were stained with 4’,6-diamidino-2-phenylindole (DAPI; 0.5 µg/mL in PBS) for 10 min at room temperature and mounted in ProLong Gold (Invitrogen).

### Confocal Microscopy

Confocal microscopy was conducted using an inverted SP8 laser scanning confocal microscope (Leica Microsystems) equipped with a compact supply unit which integrates a LIAchroic scan head, several laser lines (405 and 488 nm), and standard photomultiplier tube detectors. We used the oil immersion lens HC PL APO 63×/1.40 OIL CS2. The scanning speed was set at 400 Hz unidirectional. The pinhole was adjusted to one Airy unit for all channels. The spatial sampling rate was optimized according to Niquist criteria, generating a 0.058 × 0.058 × 0.299-µm voxel size (xyz). The Z-stack height fitted the specimen thickness. A two-step sequential acquisition was designed to collect the signal from three or four channels. The first step recorded the anti-tubulin fluorescence signal (excitation, 488 nm; emission, 530 nm) and the transmitted light. The second step acquired the DAPI fluorescence signal (excitation, 405 nm; emission, 415–480 nm). Signal intensity was averaged three times. The Fiji software was used to optimize the raw images, including maximum intensity projection and de-noising (3*3 median filter). For any given data, both wild-type and mutant images were analysed simultaneously with similar settings.

Tracking of microtubule bundles was performed on maximum intensity projections of z-planes covering the whole thickness of the cells. We drew a line transversely, perpendicular to the growth axis of the cell and crossing the nucleus. Peaks corresponding to the microtubule bundles were then identified in plots of intensity profiles and counted, in order to estimate the number of microtubule bundles in each cell. Note that in the *bas* mutants, due to the disorganized nature of the microtubule network, average bundle numbers may be somewhat underestimated. This is because this method is well adapted for tracking microtubule bundles oriented with their long axis parallel to the image plane, but we may have missed bundles that were perpendicular to the plane of the transection.

### TEM and FIB-SEM

For transmission electron microscopy (TEM) and focused ion beam-scanning electron microscopy (FIB-SEM), freshly released gametes were collected in cellulose capillaries and cultivated in the capillaries in Provasoli-enriched seawater for 3 to 5 days at 14°C in the dark or approximately 1 day at 14°C in 12 h light/12 h dark regime to produce two- to five-cell stage partheno-sporophytes. Cells in capillaries were frozen at high-pressure (HPF Compact 03, Engineering Office M. Wohlwend GmbH), freeze-substituted (AFS2, Leica Microsystems) with 0.2% OsO4 and 0.1% uranyl acetate in acetone containing 5% H2O as substitution medium (Read et al., 2021) and embedded in Epon. Ultrathin sections were stained with uranyl acetate and lead citrate and analysed with a Tecnai Spirit (Thermo Fisher Scientific) operated at 120 kV.

In order to identify a region of the sample containing algae at high density for FIB-SEM data acquisition, a 3D X-ray tomogram of the resin block was acquired with a Bruker Skyscan 1272. The region of interest was exposed using a 90°diamond trimming knife (Cryotrim 90, Diatome) mounted on a Leica UC7 ultramicrotome. The sample was then attached on a stub using conductive silver epoxy resin (Ted Pella) and gold sputter coated (Quorum Q150RS).

FIB-SEM imaging was performed with a Zeiss Crossbeam 540 or a Crossbeam 550, using Atlas 3D (FIBICS, Carl Zeiss Microscopy) for sample preparation and acquisition. After the deposition of a protecting platinum coat on the surface above the region of interest, a 60 µm wide trench was opened to identify and image several cells in parallel. During the stack acquisition, FIB slicing was done with at 30 kV and 700 pA current. The datasets were acquired at 8 nm isotropic voxel size with the SEM at 1.5 kV and 700 pA current, using a back-scattered electron detector (ESB). After acquisition, the image stacks were acquired using the Fiji plugin “Linear stack alignment with SIFT” (Lowe, 2004). We acquired images for a total of 5 cells for wild-type and 5 *bas-2* cells, and chose a representative image to present in Figure 4. Images containing Golgi stacks were retrieved from aligned image stacks and cropped to reduce the image dimensions for further segmentation with the IMOD software package (https://bio3d.colorado.edu/imod/).

### Phylogenetic trees

Multiple alignments were generated with Muscle in MEGA7 (Kumar et al., 2016). Phylogenetic trees were then generated with RAxML (Stamatakis, 2014) using 1000 bootstrap replicates and the most appropriate model.

### *BAS* and *DIS* gene expression estimation during the *Ectocarpus* life cycle

Expression levels of the *BAS* and *DIS* genes were investigated using TPM values obtained after kallisto pseudo-mapping and calculation of the lengthScaledTPM using the tximport package in R (see ‘Comparative Transcriptome Analyses’ section for details). Previously generated RNA-seq data for wild type and *dis* samples (Table S7) was used for comparisons.

### Comparative transcriptome analyses

RNA-seq analysis was performed to compare the abundances of gene transcripts in the mutants *bas-1* and *bas-2* and wild-type sporophytes. RNA-seq datasets were generated from triplicate samples of each genotype, and individuals were grown synchronously as described previously in standard culture conditions (Coelho et al., 2012b; Cossard et al., 2022). Germlings were filtered through a nylon mesh to recover only thalli at the 2-5 cell stage. Each replicate contained between 10^4^ and 10^6^ individual germlings. Tissue samples were rapidly frozen in liquid nitrogen and processed for RNA extraction. Total RNA was extracted from each sample using the Qiagen Mini kit as previously described (Lipinska et al., 2015). For each replicate, cDNA was produced by oligo(dT) priming, fragmented, and prepared for stranded 2× 150-bp paired-end sequencing on an Illumina HiSeq 3000 platform.

Raw and cleaned read quality was evaluated using fastQC (v. 0.11.9) and mutliQC (v. 1.9). Raw reads were trimmed and filtered based on quality score, and adapter sequences were removed using Trimmomatic (v. 0.39). Transcript abundance was evaluate using kallisto (v. 0.46.2) via pseudo mapping on CDS features. Then matrix of counts and TPMs for all samples, all replicates were generated in R using tximport package.

A gene was considered to be expressed if the TPMmean was above the 5^th^ percentile (as in Cossard et al., 2022; Lipinska et al., 2019). About 4% of the genes in each sample had TPMmean values under this threshold and were considered not to be expressed in our samples (Table S9). Differential gene expression was analysed using DESeq2 (Love et al., 2014, 2). Genes were considered to be differentially expressed when the log2 fold change was below or equal −1 (down-regulated, at least twice as weakly expressed) or above or equal to 1 (up-regulated, at least twice as strongly expressed) and adjusted p-value below or equal to 0.01. Genes were considered to be exclusively (uniquely) expressed in *bas* mutants when they were significantly up-regulated compared to wild-type and the wild-type mean TPM was equal to 0. Conversely, genes were considered to be specifically silenced in *bas* mutants when there were significantly down-regulated compared to wild-type and their TPM means were equal to 0.

GO term and HECTAR localisation enrichment were carried out on R using the “clusterProfiler” package (Yu et al., 2012).

## Accession Numbers

Accession numbers are provided in Table S7.

## Acknowledgements

This work was supported by the CNRS, Sorbonne Université, the Max Planck Society and the European Research Council (grant agreement 864038 to SMC). HY was supported by a grant from the China Scholarship Council (grant number 201608310119). We thank Nana Kinoshita-Terauchi (Shimoda Marine Research Center, University of Tsukuba) and Iris Kock (MPI Tübingen) for helpful discussion and images of the mutants and Viktoria Buhanets for help with segmentation of Fib-Sem images.

## Author contribution

S.M.C, J.M.C and O.G conceived the project. O.G., M.Z. and S.M.C. performed the main experimental work. O.G. performed bioinformatic analyses. J.M.C. carried out gene annotation and phylogenetic analyses. Y.H., M.Z, A.H, A.F.P performed experiments. S.C. and D.S. generated epifluorescence images. P.R. and K.H. generated and analysed Fib-Sem data. C.N and T.M generated TEM images. O.G., J.M.C. and S.M.C. interpreted the data. S.M.C. wrote the manuscript with input from all authors.

## Supplemental Figures

**Figure S1.**
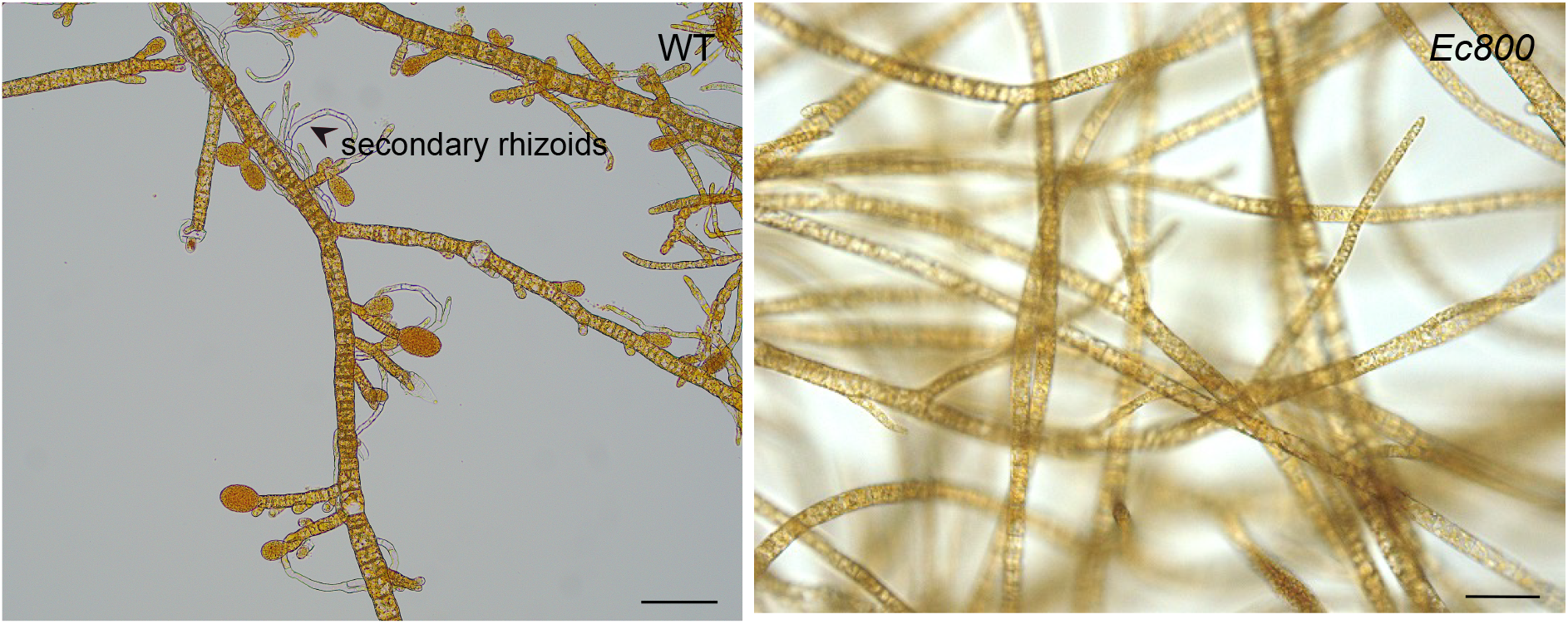
Absence of secondary rhizoids in apical filaments of mutant lines compared with wild-type, after 3-weeks in culture. Scale=20 µm.

**Figure S2.**
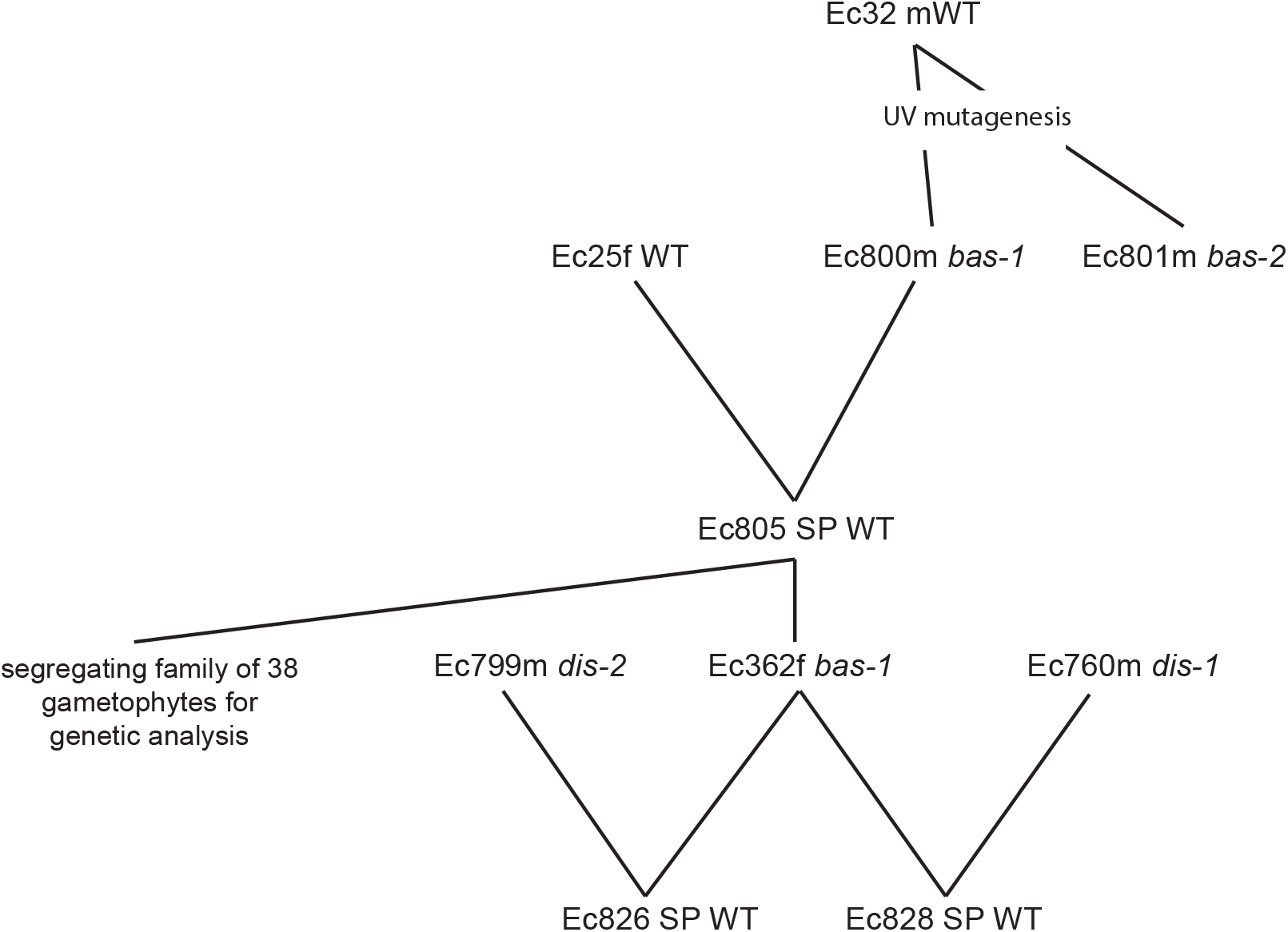
Pedigree of the *Ectocarpus* strains used in this study. SP, diploid, hybrid sporophyte; WT, wild type; m, male; f, female.

**Figure S3.**
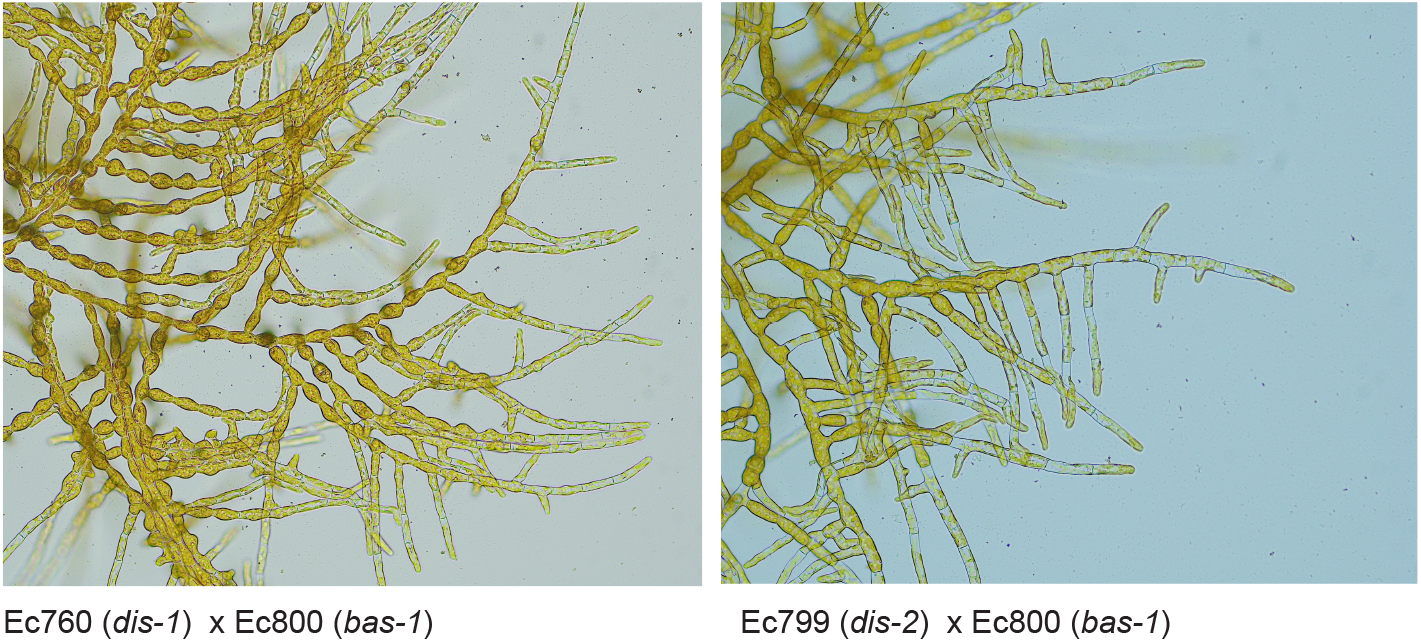
Morphological phenotypes of diploid sporophytes derived by crossing *dis-2* x *bas-1* or *dis-1* x *bas-2*. Note that the diploid sporophyte derived from the cross has a wild-type phenotype, indicating the two mutations complement each other. Scale=20 µm.

**Figure S4.**
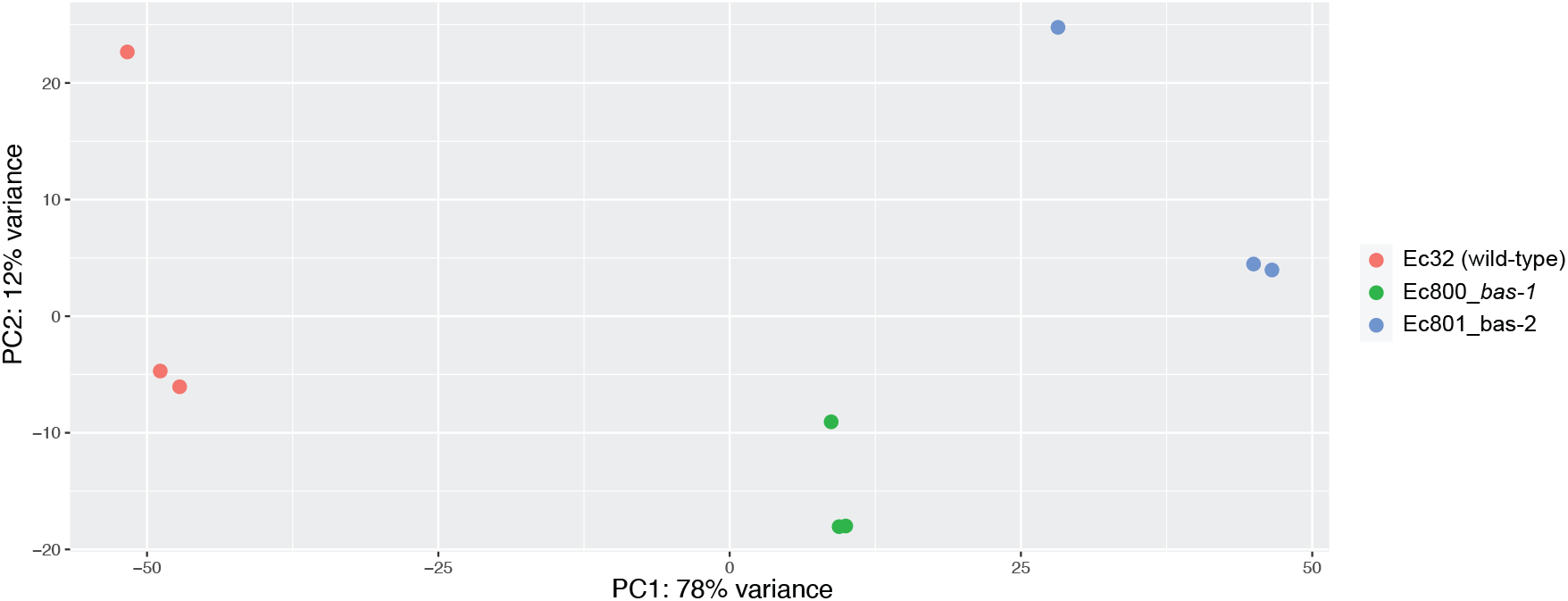
Principal component analysis (PCA) comparison of transcript abundance patterns for all expressed genes across wild-type (WT), *bas-1* and *bas-2* replicate samples. The two dimensions represent 78% and 12% of the variance. The analysis was carried out using normalized counts generated by DESeq2 after Variance Stabilizing Transformation (VST).

**Figure S5.**
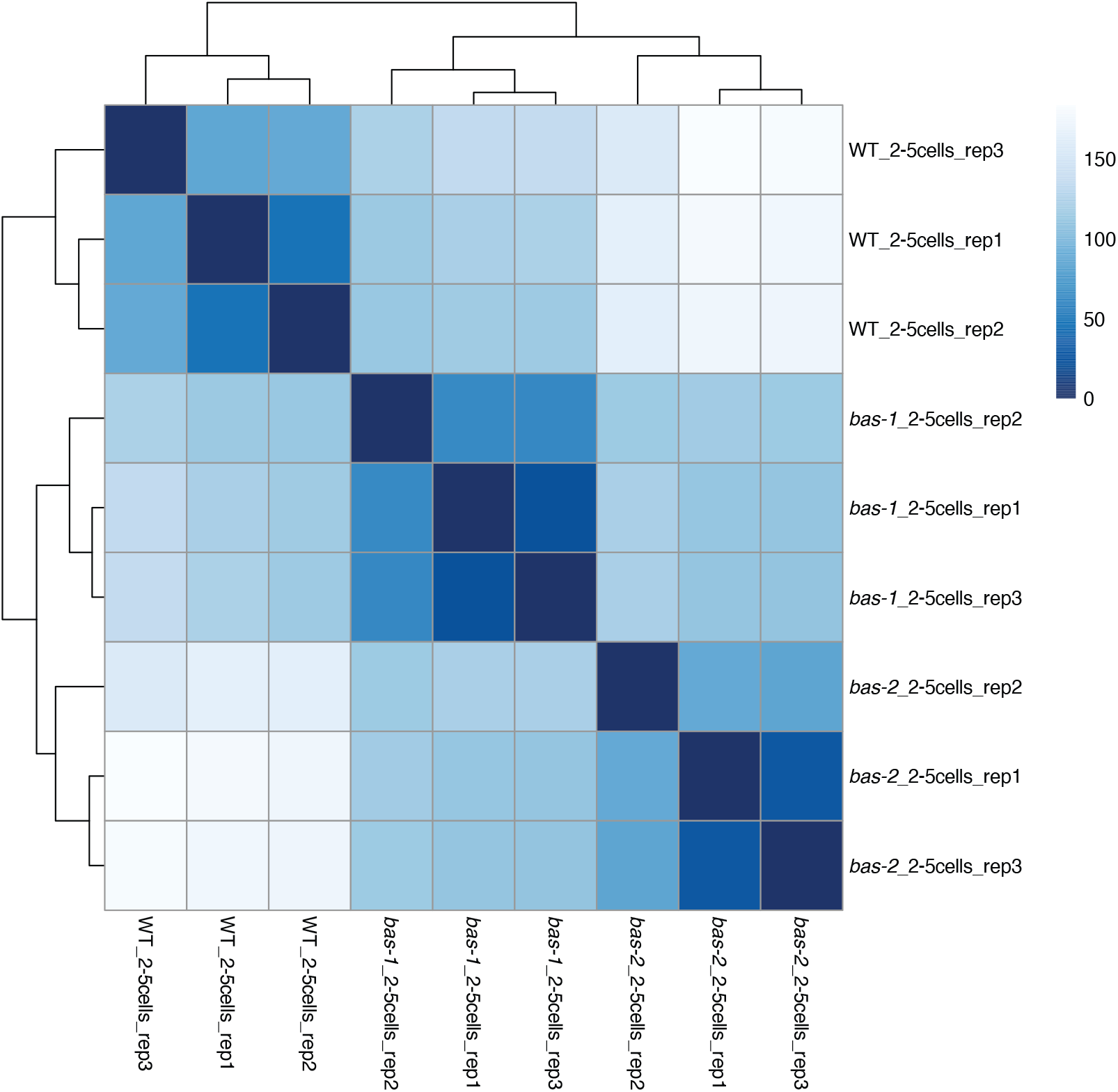
Between-sample correlation diagnostics of the RNA sequencing data. Heat map representing the distance between replicates. Euclidian distances were calculating from normalized counts generated by DESeq2 after Variance Stabilizing Transformation (VST). Details of the samples are given in Table S7.

**Figure S6.**
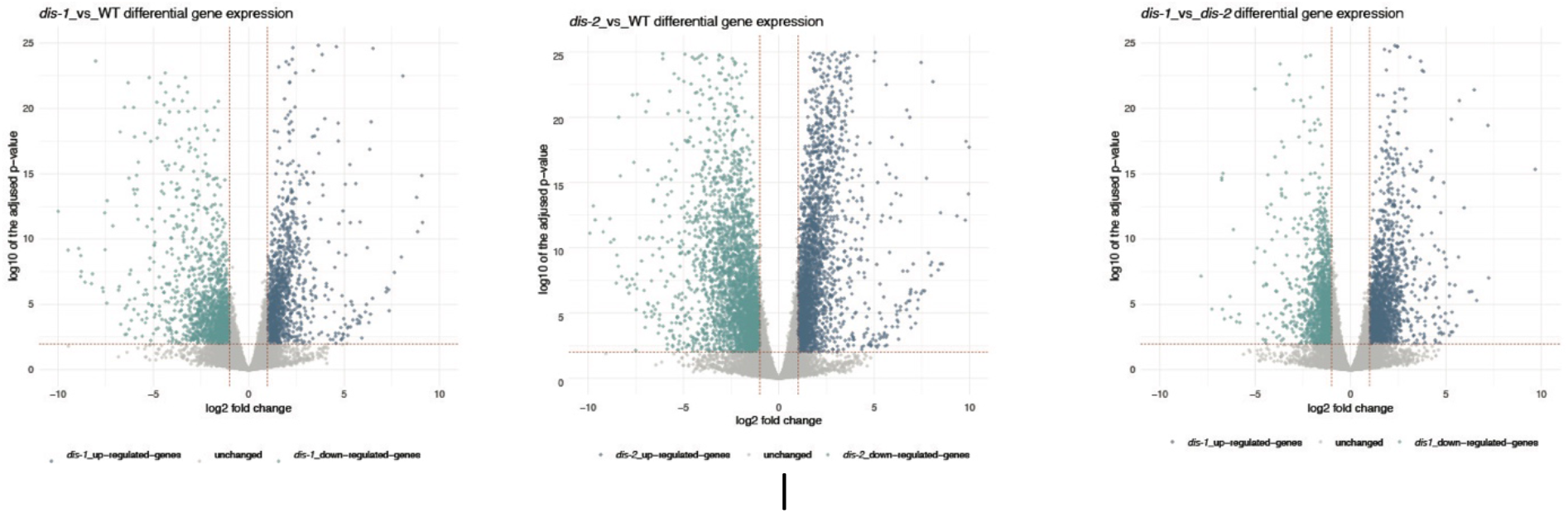
Vulcano plots of all genes in pairwise comparisons between wild-type (WT) and mutants (*bas-1* and *bas-2*). The log2 FC value was calculated based on the mean expression level (TPM) for each gene. Each dot represents one gene. Blue represents upregulated genes and green downregulated genes in each comparison.

**Figure S7.**
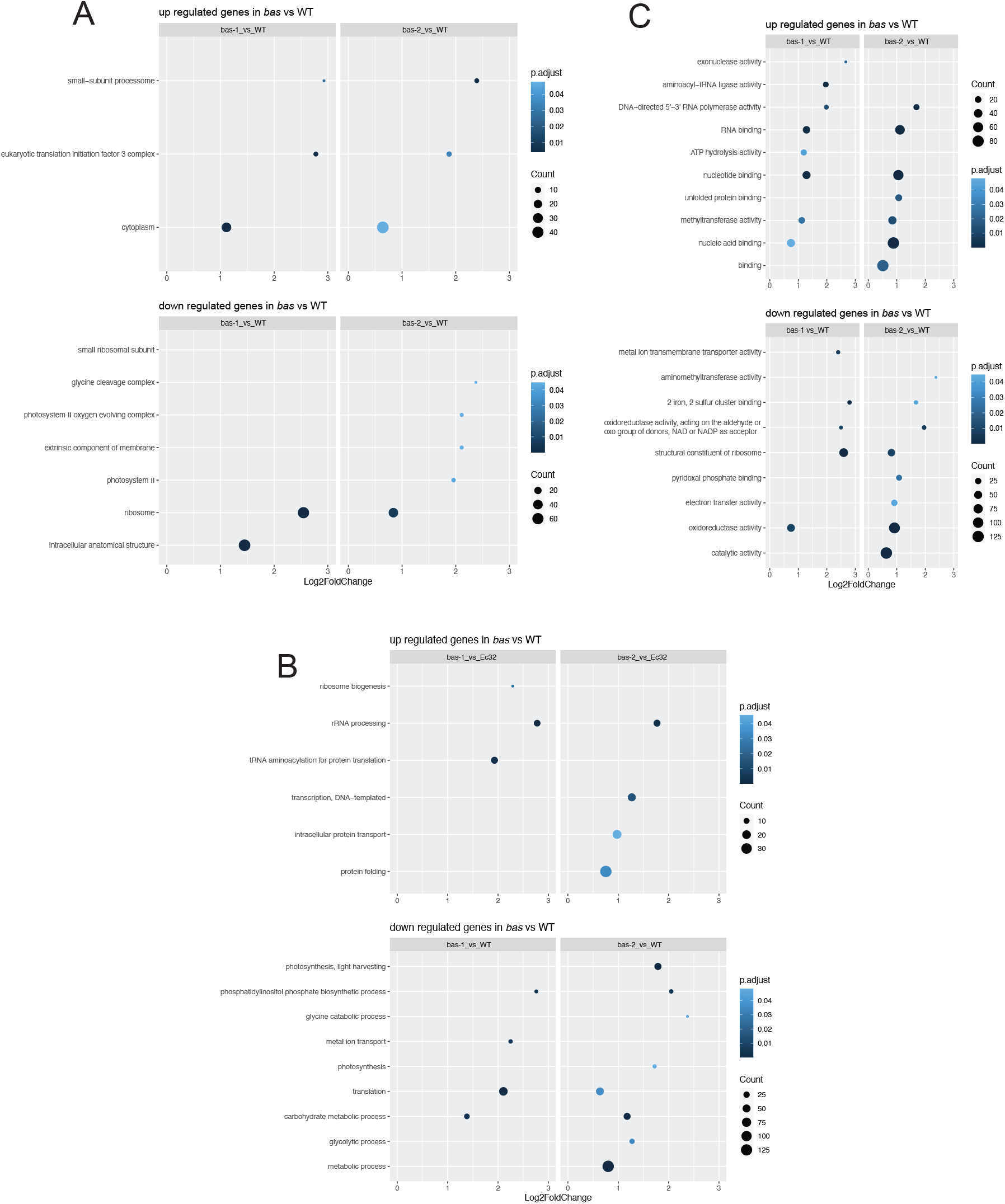
GO term enrichment observed in DE gene sets in bas mutants compared to WT. Dot plot representation is divided according to GO term ontology classes ‘Cellular Component’ (A), ‘Molecular Function’ (B) and ‘Biological Processes’ (C).

**Figure S8.**
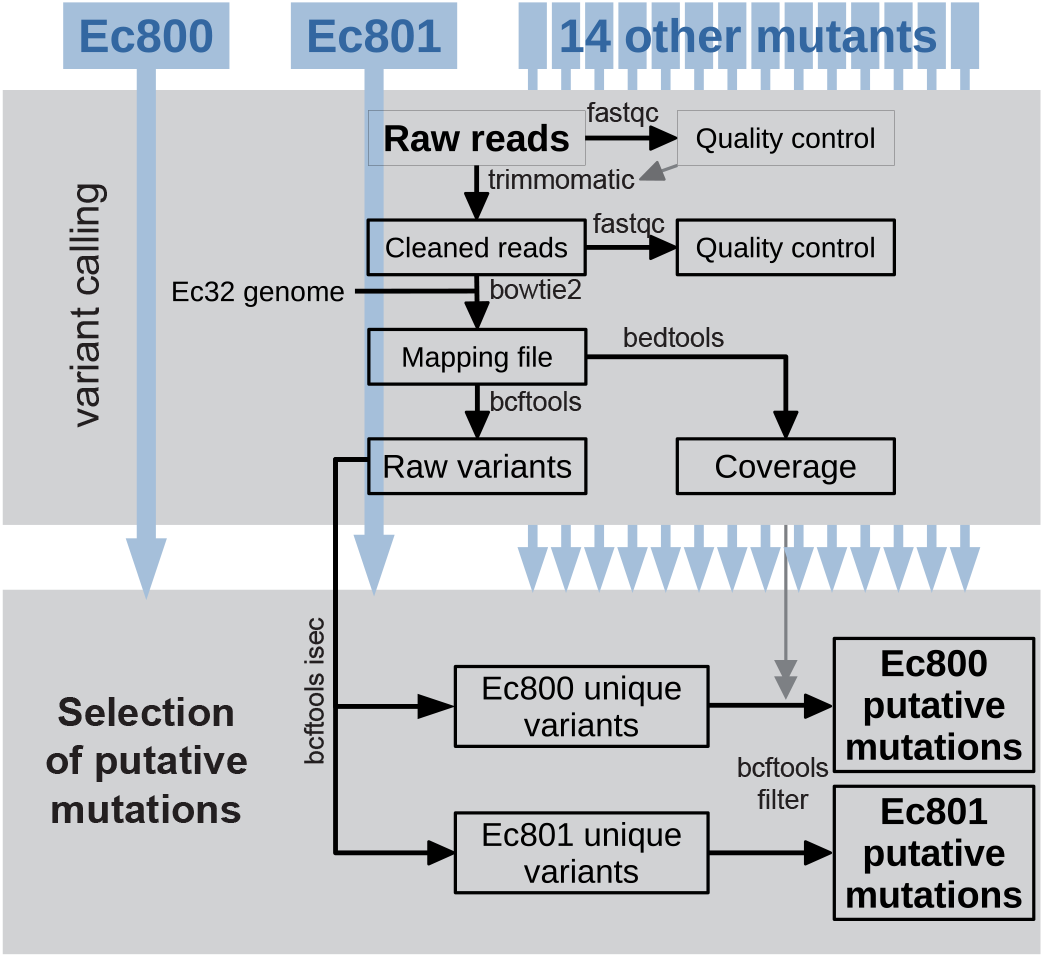
Schematic diagram of the approach used to detect putative mutations in the genomes of Ec800 and Ec801.

